# The language of music: Common neural codes for structured sequences in music and natural language

**DOI:** 10.1101/202382

**Authors:** Jeffrey N. Chiang, Matthew H. Rosenberg, Carolyn A. Bufford, Daniel Stephens, Antonio Lysy, Martin M. Monti

## Abstract

The ability to process structured sequences is a central feature of natural language but also characterizes many other domains of human cognition. In this fMRI study, we measured brain metabolic response in musicians as they generated structured and non-structured sequences in language and music. We employed a univariate and multivariate cross-classification approach to provide evidence that a common neural code underlies the production of structured sequences across the two domains. Crucially, the common substrate includes Broca’s area, a region well known for processing structured sequences in language. These findings have several implications. First, they directly support the hypothesis that language and music share syntactic integration mechanisms. Second, they show that Broca’s area is capable of operating supramodally across these two domains. Finally, these results dismiss the recent hypothesis that domain general processes or proximal neural substrates explain the previously observed “overlap” between neuroimaging activations across the two domains.

---

The language of music: Common neural codes for structured sequences in language and music A central intuition in the study of human language as a cognitive phenomenon is the idea that, while listening to a linear signal such as speech, our minds spontaneously build abstract and structured hypotheses representing how discrete elements within a sequence relate to each other (Monti, 2017). The use of such representations is most clearly displayed in natural language, but also characterizes other aspects of human cognition, such as logic reasoning (Osherson, 1975; Monti & Osherson, 2012), algebraic cognition (Varley et al., 2005; Monti et al., 2012; Maruyama et al., 2012), and music cognition (Patel, 2003; Katz & Pesetsky, 2011; Lerdahl, 2001), among others. The relationship between the syntactic operation of language and the syntax-like operations of other aspects of human cognition has thus been the at the center of a long-standing debate concerning the degree to which human thought is embedded within, or enabled by, natural language (e.g., Lashley, 1951; Boeckx, 2010; Gleitman & Papafragou, 2013; Fitch & Martins, 2014; Fitch 2014; Monti, 2017).

Lashley (1951) commented on the prevalence of structured sequences across domains, noticing that they exhibited the following three properties: (1) connectedness; i.e. no node is isolated from the others, (2) a root element; i.e. “sentence” or “chord” that is superior to others and (3) acyclic structure; establishing order as a unique property (Lashley 1951, Fitch and Martins 2014). In the context of music cognition, the analogy with the structural aspects of language is particularly pronounced. As discussed elsewhere (e.g., Lerdahl & Jackendoff, 1985; Patel, 2003; Fadiga et al., 2009; Fitch 2014; Peretz et al., 2015), music and language are both characterized by discrete elements (e.g., words, chords) which can be (recursively) combined, according to specific rules, to form organized structures (e.g., sentences, melodies) which are typically encoded within linear, time-dependent, signals.

Nonetheless, whether this analogy is substantial or merely superficial remains a debated issue (cf., Peretz et al., 2015). At one end of the spectrum, it has been proposed that language and music are governed by the very same syntactic processes applied to different building blocks (e.g., words vs. notes). According to this view, “[a]ll formal differences between language and music are a consequence of differences in their fundamental building blocks[; i]n all other respects, language and music are identical” (Katz & Pesetsky, 2011). Along similar lines, it has been proposed that the common representations underlying the structure processing in language and music can be localized to the neural mechanisms encapsulated within the left inferior frontal gyrus (IFG; often referred to as Broca’s Area), a region hypothesized to operate as a “supramodal hierarchical parser” (Tettamanti & Weniger, 2006; Fadiga et al., 2009). Consistent with this view, a rapidly growing neuroimaging literature has shown music processing to recruit cortical regions overlapping with areas known to be involved in syntactic and semantic aspects of natural language processing (Patel et al., 1998; Maess et al., 2001; Koelsch et al., 2002; Tillmann et al., 2003; Koelsch et al., 2004; Koelsch et al., 2005; Brown et al., 2006; see Rogalsky et al., 2011, for a conflicting result). Nonetheless, while the observation of overlapping neural substrates is often taken to imply the presence of shared neurocognitive representations between language and music, this is not necessarily the case (Peretz et al., 2015) and indeed has never been shown to be true. This “missing link” in the neuroscientific literature leaves open the possibility that commonly recruited areas of the brain might, in fact, represent very different operations that do not translate, or align, across the two domains, or that are entirely unrelated to the processing of these relationships. In line with this observation, it has been suggested that language and music are in fact better thought of as modular and largely independent of each other (Marin & Perry, 1999; Peretz & Coltheart, 2003). In support of this view, a rich neuropsychological literature has described cases of individuals who exhibit amusia in the absence of aphasia, as well as aphasia in the absence of amusia (Luria et al., 1965; Peretz, 1993; Peretz et al., 1994; Ayotte et al., 2000; Piccirilli et al., 2000; Ayotte et al., 2002).

The reason for the contradicting evidence is still a matter of debate. According to some, the fracture between neuropsychological and neuroimaging findings can be reconciled with a middle-ground solution in which language and music are viewed as partially overlapping systems (Patel, 2003; Patel et al., 2008). Under this view, referred to as the shared syntactic integration resource hypothesis, language and music are characterized by both domain-specific (i.e., separate) and domain-general (i.e., shared) processes. The domain-specific processes relate to the particular features of each syntax, which are recognized as architecturally different, while domain-general processes provide neural resources for the activation of the relevant stored syntactic representations (Patel, 2012). According to others, the inconsistency between the two sets of findings might instead be due to experimental and neuroanatomical considerations (Fedorenko & Varley, 2016). Specifically, the overlap often reported, in neuroimaging studies, in left inferior frontal regions could be a reflection of task-general demands tied to the use of structural-violation paradigms (e.g., the P600 and the early left/right anterior negativity effects reported in electrophysiological studies; Janata, 1995; Maess et al., 2001; Koelsch et al., 2002, 2005; Steinbeis and Koelsch, 2008; Tillmann et al., 2003; and later localized to the inferior frontal gyri through neuroimaging; Musso et al 2015; Kunert et al 2015). Deviant events are indeed likely to elicit ancillary processes including attentional capture, detection of violated expectations, or error correction, regardless of whether the violation applies to natural language, music, arithmetic, or motor sequences. Such processes are unrelated to the extracting or forging of structured sequences and are known to elicit activation in domain-general regions (proximal or partially overlapping with Broca’s Area; see Fedorenko & Varley, 2016, for a detailed discussion).

In the present study, we address the relationship between the mechanisms of natural language and those of music in a 3 Tesla functional magnetic resonance imaging (fMRI) within-subjects design in which competent musicians generate structures in language (active/passive voice sentences versus repeating a verb) and music (root/second-inversion position ascending triads versus repeating a note; cf., Figure 1 and Table 1). Crucially, we employ a (rarely explored) generation task to avoid the confound of salient events, and we use a multivariate cross-classification approach to resolve the interpretational ambiguity present in the previous neuroimaging literature (which has been specifically advocated for; see Peretz et al., 2015), thereby helping resolve the question of whether natural language and music share a common underlying neural code for representing structured sequences.

## Methods

### Participants

We recruited 21 total participants to reach the predetermined sample size (N=20, 8 female participants) based on previous literature (Musso et al, 2015: N=11; Kunert et al, 2016: N=19; Koelsch et al, 2002: N=20). An additional subject was recruited because the data from one of the participants exhibited excessive motion during the procedure (see below). Participants received $50 compensation for taking part in the experiment. All participants were native English speakers, right handed, and competent musicians currently enrolled in the UCLA Herb Alpert School of Music. Participants were only enrolled if they could demonstrate proficiency in singing/generating both a root position and II^nd^ inversion ascending triad arpeggio. Participants with perfect pitch were excluded. Participants signed informed consent prior to taking part in the session, as per the procedures approved by the UCLA Institutional Review Board.

### Stimuli

For both materials (i.e., “language” and “music” trials), the first cue was delivered visually, by presenting one of three icons in the middle of the screen. A ‘◊’ symbol indicated an active or root position trial (depending on whether the second cue was a word or a note, respectively); a ‘♣’ symbol indicated a passive or a II^nd^ inversion trial; a ‘**↺**’ symbol indicated a non-structured (i.e., repeat) trial. The second cue was delivered aurally and consisted of either a verb or a note, thus revealing whether the trial was a language or music trial, and allowing disambiguation of the instruction provided by the first cue. (See Table 1 and Figure 1 for sample stimuli.) Cues for language trials consisted of seven monosyllabic, reversible, present tense verbs (i.e., “bring,” “tell,” “teach,” “throw,” “leave,” “give,” “pay”). Cues for the music trials consisted of 7 notes recorded with an electronic piano. The notes comprised a chromatic scale (i.e. each consecutive pitch was separated from the following by a semitone, or a half step). Participants were allowed to choose, among three sets of musical cues (high, middle, or low), whichever pitch range most comfortably matched their vocal range. For each set, a perfect fifth (7 semitones/half steps) separated the lowest and highest pitches of the cues. Combination of the 7 cues (i.e., words/notes), 3 generative rules (2 structured, 1 repeat), and 2 materials (i.e., language, music), resulted in 42 unique trials (i.e., 21 per material type).

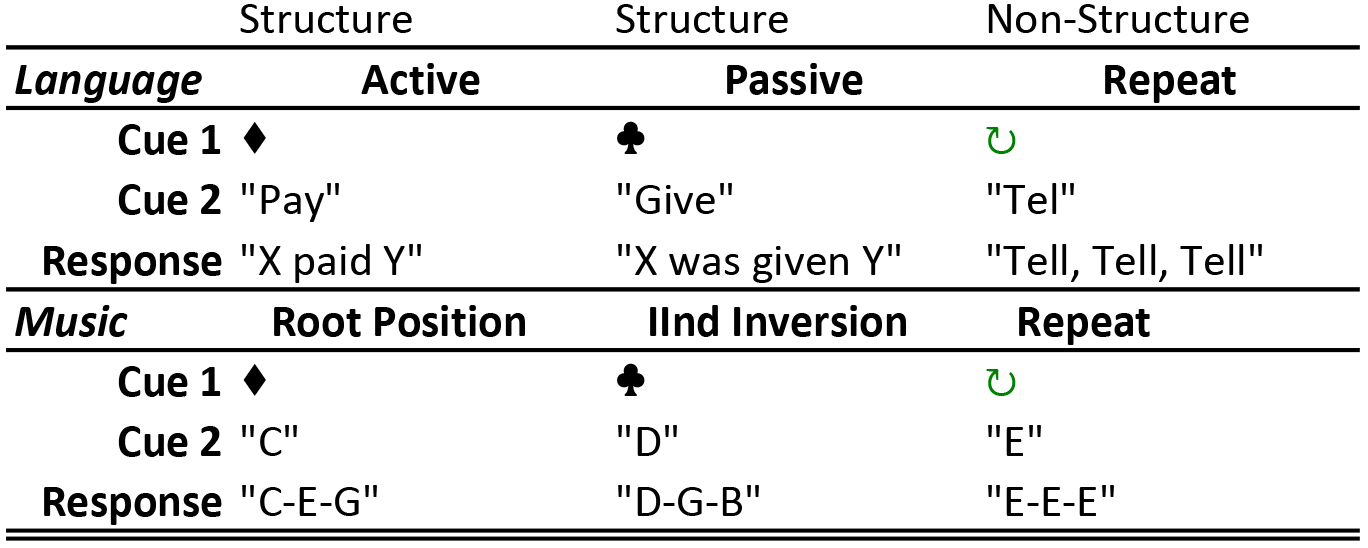

### Experimental Design

Each participant viewed the 42 unique trials twice (once in the first two runs, once in the second two runs). Trial types were equally distributed across 4 runs, and, within each, randomly presented. Stimuli were presented using PsychoPy (Peirce, 2008); visual cues were displayed through a custom-made MRI-compatible projection system while auditory cues were delivered through a Magnetic Resonance headphone system. As shown in Figure 1, each trial started with the generative rule cue (i.e., ‘◊,**♣**,**↺**’), displayed on screen for 1.5 s, followed by the second cue (i.e., word or note) presented, aurally, for 1.8 s. After a variable jitter (between 6 and 8 s), a fixation symbol blinked four times (with a cycle of 0.8 s of display and 0.35 s interval). The first blink (with a black square symbol) served as a warning that the “performance/response” period was to begin. The following three blinks (with a black circle symbol) marked the performance/response period and provided a tempo for responding. The tempo was never varied, neither within nor across subjects, and was only employed to provide participants with a consistent rhythm for responding.

**Figure 1.**
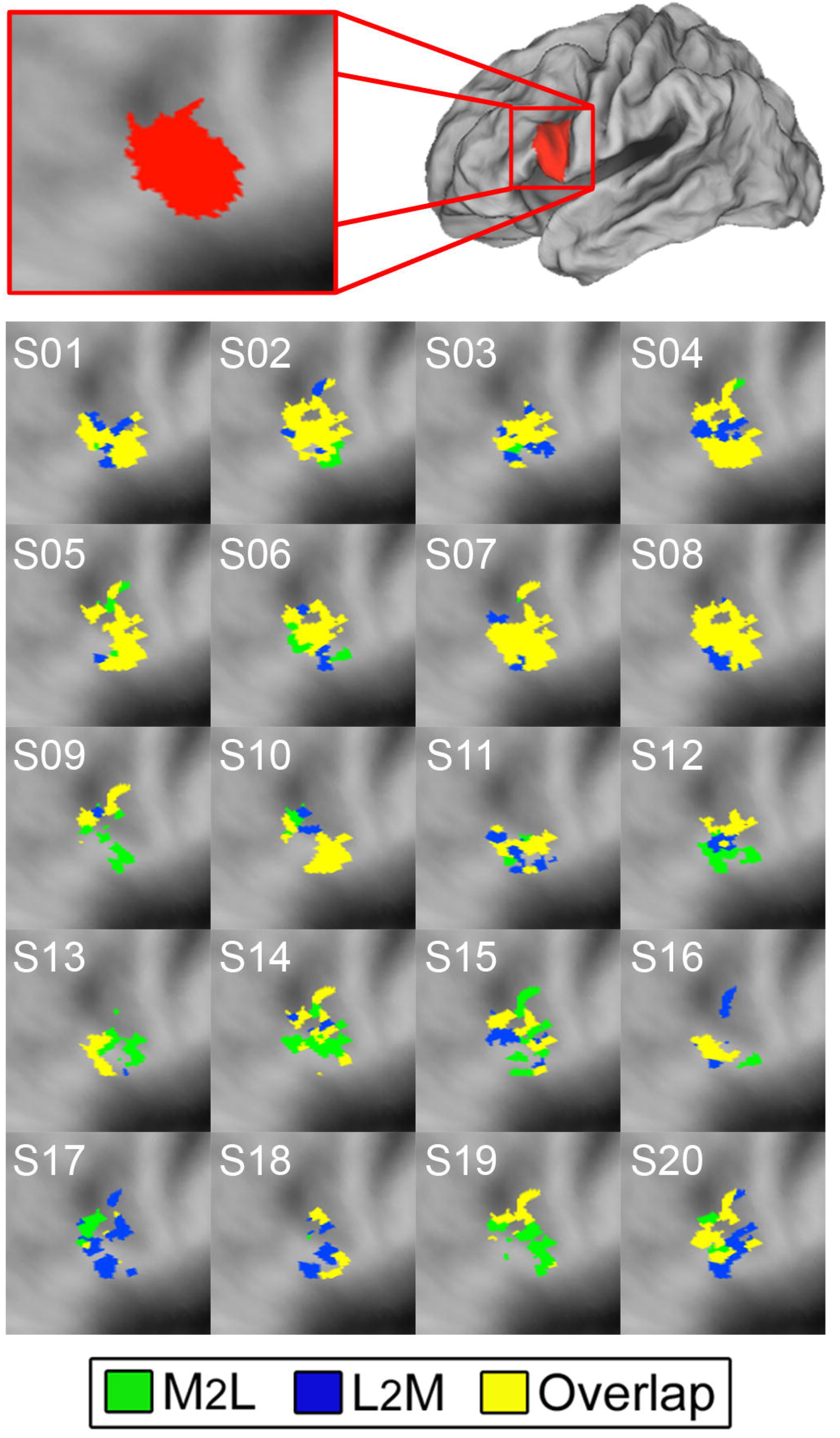
Experimental design. Sample music and language trials timelines.

Finally, a variable length fixation screen (with a random jitter between 5 and 7 s chosen, on a trial-by-trial basis, from an exponential distribution) separated each trial from the subsequent one. Each run lasted, on average, 293.57 s (S.D. = 15.81). Participants were trained to asymptotic performance prior to the imaging session, in a separate room, after having signed informed consent. The experimenter corrected any errors the participant made until satisfactory performance was achieved (less than 2 errors per block of trials). Training ceased when participants could perform at least 12 out of 13 trials correctly, minimizing the sound production time across conditions.

### Data Acquisition

Data were acquired on a 3 Tesla Siemens Tim Trio Magnetic Resonance Imaging (MRI) scanner at the Staglin IMHRO Center for Cognitive Neuroscience at UCLA. Structural data were acquired using a T1-weighted sequence (MP RAGE, TR = 1,900 ms, TE = 2.26 ms, voxel size 1 mm^3^ isovoxel). Blood oxygenation level dependent (BOLD) data were acquired with a T2*-weighted Gradient Recall Echo sequence (TR = 3,000 ms, TE = 35 ms, 45 interleaved slices, voxel size 3 × 3 × 3.3 mm) with prospective motion correction in order to reduce the impact of subject motion during performance.

### Data Preprocessing

Data analysis was carried out using FSL (Smith et al., 2004). Prior to analysis, data underwent a series of conventional preprocessing steps including motion correction, slice-timing correction (using Fourier-space time-series phase-shifting), spatial smoothing using a Gaussian kernel of 5 mm full-width half-max, and highpass temporal filtering (Gaussian-weighted least-squares straight line fitting, with σ=50.0s). Data from each individual run were analyzed employing a univariate general linear model approach (Monti, 2011) inclusive of a pre-whitening correction for autocorrelation. Following current convention, any participant exhibiting average motion greater than 3 mm was excluded (N=1).

### Univariate Analysis

For each run of each participant, a univariate analysis was conducted using, as the main variables of interest, 6 regressors, one per trial type (i.e., language active voice, language passive voice, language repeat, music root position, music II^*nd*^ inversion position, music repeat). Regressors marked the performance/response period of each trial (see Figure 1). A number of additional nuisance regressors were employed to model cue periods, motion (including first and second derivatives, and their difference), as well as the short intervals between the second cue and task performance. This last regressor is particularly important since it parcels out periods in which subjects are likely to be engaging in strategies in anticipation of the task, which, in the absence of any participant feedback, are un-controlled and thus difficult to interpret. For each run we computed 4 contrasts: structured versus repeat trials for language and music materials (“simple effect” contrasts), separately, and the interaction between the two simple effects (“interaction contrasts”) in both directions (i.e., simple effect of structured trials in language greater than the simple effect of structured trials in music, and vice versa). Data from individual runs were aggregated employing a mixed effects model (i.e., employing both the within- and between-subject variance), and using automatic outlier detection. Z (Gaussianised T) statistic images were thresholded using a cluster correction of Z>2.3 and a (corrected) cluster significance threshold of P=0.05.

### Multivariate Analysis

The input to the multivariate analysis was a set of volumes of regression coefficients (i.e., 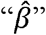) marking the magnitude of activation, for each voxel, in each trial (per subject). These trial-wise “patterns of activations” were obtained by employing the iterative Least Squares – Separate approach (LS-S; Mumford et al., 2012) in which a separate GLM is run (here, using FILM with local autocorrelation) for each trial. At each iteration, one regressor marks the trial of interest, while all remaining trials are collapsed into a nuisance regressor (see Mumford et al., 2012, Figure 1 for a visual depiction of this approach). This approach has been shown, in simulations, to produce activation estimates that have the highest correlation with true activation magnitudes (Mumford et al., 2012), and has also been shown to adapt best to multivariate analyses when used in conjunction with full randomization of trials (different for each subject) and with equal inter stimulus interval across condition (Mumford et al., 2014), as we have done. The patterns of activation were then concatenated across time to construct a subjectwise “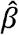-series” of activation magnitude per trial per voxel (Rissman et al., 2004). In order to assess whether natural language and music share underlying neural representations, we employed a cross-classification searchlight analysis using a linear support vector machine (SVM) algorithm. Cross-classification was performed by training the SVM classifier to recognize structure vs. repeat structure trials in one domain, and then attempting to classify structure vs. repeat structure trials in the other domain (“M2L” and “L2M” cross-classifications for training on music and testing on language and vice versa, respectively). L2M cross-classifications were performed over voxels found significant in the structure minus repeat trials for language materials (only); M2L classifications were performed over voxels found significant in the structure minus repeat trials for music materials (only). While the significant voxels in the two univariate contrasts could overlap, this feature selection ensures that the training and testing datasets for each type of cross-classification (i.e., L2M, M2L) remain completely separate, thereby avoiding any bias in the analysis. Classifications were performed on a single subject basis, in native space, employing a 6 mm radius searchlight approach (Kriegeskorte et al., 2006). To account for the imbalance between the number of structure and repeat trials (28 and 14, respectively, per each domain) and avoid biasing the classifier, we performed a resampling procedure in which, at each of 1000 iterations, a subsample of 14 (structured) trials was randomly selected, in order to train and test the classifier on a matching number of trials across conditions. Results across the 1000 iterations were averaged to yield a single classification accuracy value for each searchlight sphere. Statistical significance was assessed, at the group level, employing a permutation-based sign test and against a criterion of p= 0.05 corrected for multiple comparisons at the cluster level (using FSL’s threshold free cluster enhancement). At the single subject level, significance was assessed with the same resampling procedure, repeated 1000 times, with shuffled testing labels, to construct a null distribution for each voxel (cf., Etzel & Braver, 2013). Classifications falling within the top 5% of the null distribution were considered significant.

## Results

### Univariate Analysis

The simple effect contrast of structure versus repeat trials for language materials uncovered a set of expected activations in left inferior frontal gyrus (including its *pars opercularis* and *triangularis*, in Brodmann Areas [BA] 44, 45), posterior middle and superior temporal cortices (BA 21, 22), bilateral parietal (spanning BA 7, 40) and medial (BA 6), middle (BA 8), and superior frontal (BA 6) areas (mostly left lateralized; see Figure 2, below, and Table S1 in the Supplemental Material available online for complete list of local maxima).

**Figure 2.**
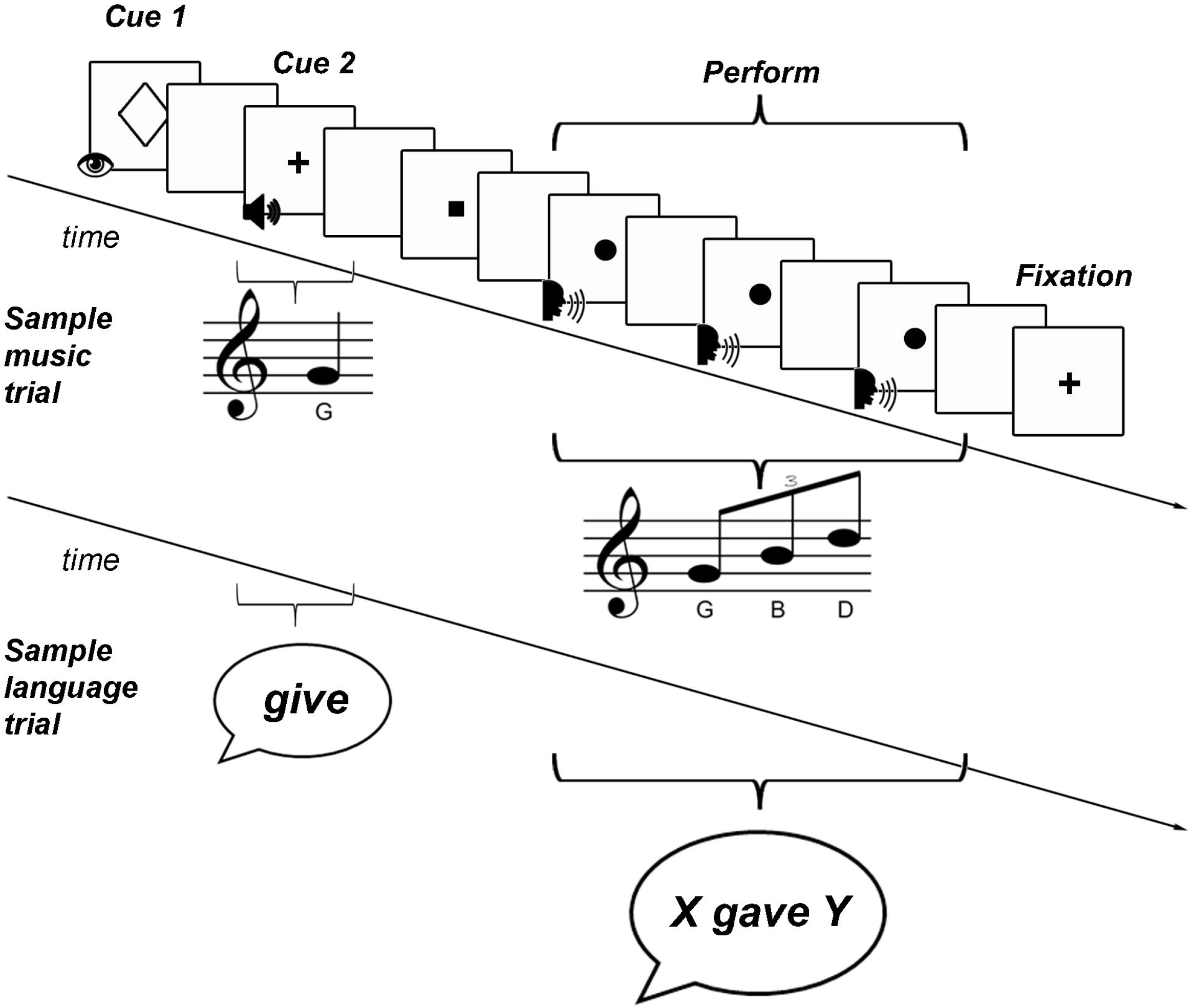
Univariate result. Overlay of the structure versus repeat contrast results for language (blue) and music (green) (yellow marks overlap between the two tasks).

When performed on music trials, the same contrast uncovered a number of activation clusters across bilateral frontal and parietal regions (see Figure 2 and Table S2). The frontal cluster included bilateral maxima in the inferior frontal gyri (spanning its *pars opercularis* in BA 44, *triangularis* in BA 45, and *orbitalis* in BA 47), rostral insular cortex (spanning BA 13 and its junction with 45 and 47), as well as bilateral foci across middle (BA 6), superior (BA 6, 8) frontal, and cingulate (BA 32) gyri. In addition, bilateral activations were observed in the inferior (BA 40) and superior (BA 7) parietal lobuli, as well as in the posterior cerebellum (see Table S2 in the Supplemental Material available online for the complete list of local maxima). As shown in Figure 2 (regions in yellow), the structure versus repeat contrast uncovered a number of common areas across language and music materials, including the left inferior frontal (in its *pars opercularis*, BA 44) and middle frontal (in BA 6) gyri, as well as the medial frontal/cingulate gyri (BA 6, 32), and bilateral posterior parietal lobe (in both BA 7 and 40). In order to avoid interpreting a “reverse subtraction”, we characterized the mean activity profile for structure and repeat conditions to identify the primary driver in IFG. Mean zscores from the IFG subregions (defined by external atlases: pars opercularis and pars triangularis from Harvard-Oxford and pars orbitalis from AAL) are displayed in figure S2.

The interaction of structure versus repeat structure and materials revealed the left superior and middle temporal gyri (BA 21, 22) to be specific to language (see blue areas in Figure S1 and Table S3), whereas foci surrounding the right orbital and sub-lobar segments of the inferior frontal gyrus (mainly in BA 47 and 13), along with right superior frontal (BA 6), medial frontal (BA 6, 32) and contralateral posterior cerebellum appeared to be specific to structure in music (see Figure S1 and Table S3 in the Supplemental Material available online).

### Multivariate Analysis

In order to assess whether natural language and music share neural codes for representing structure, we performed a cross-classification multivariate pattern analysis. In this approach, a support vector machine (SVM) classifier was trained to recognize structure versus repeat trials in one domain and was then tested on the other. That is to say, we trained a classifier to distinguish structure versus repeat trials in language and then tested it by assessing its ability to discriminate structure versus repeat trials in music (“L2M” cross-classification), and vice versa (“M2L” cross-classification; see Methods).

**Figure 3.**
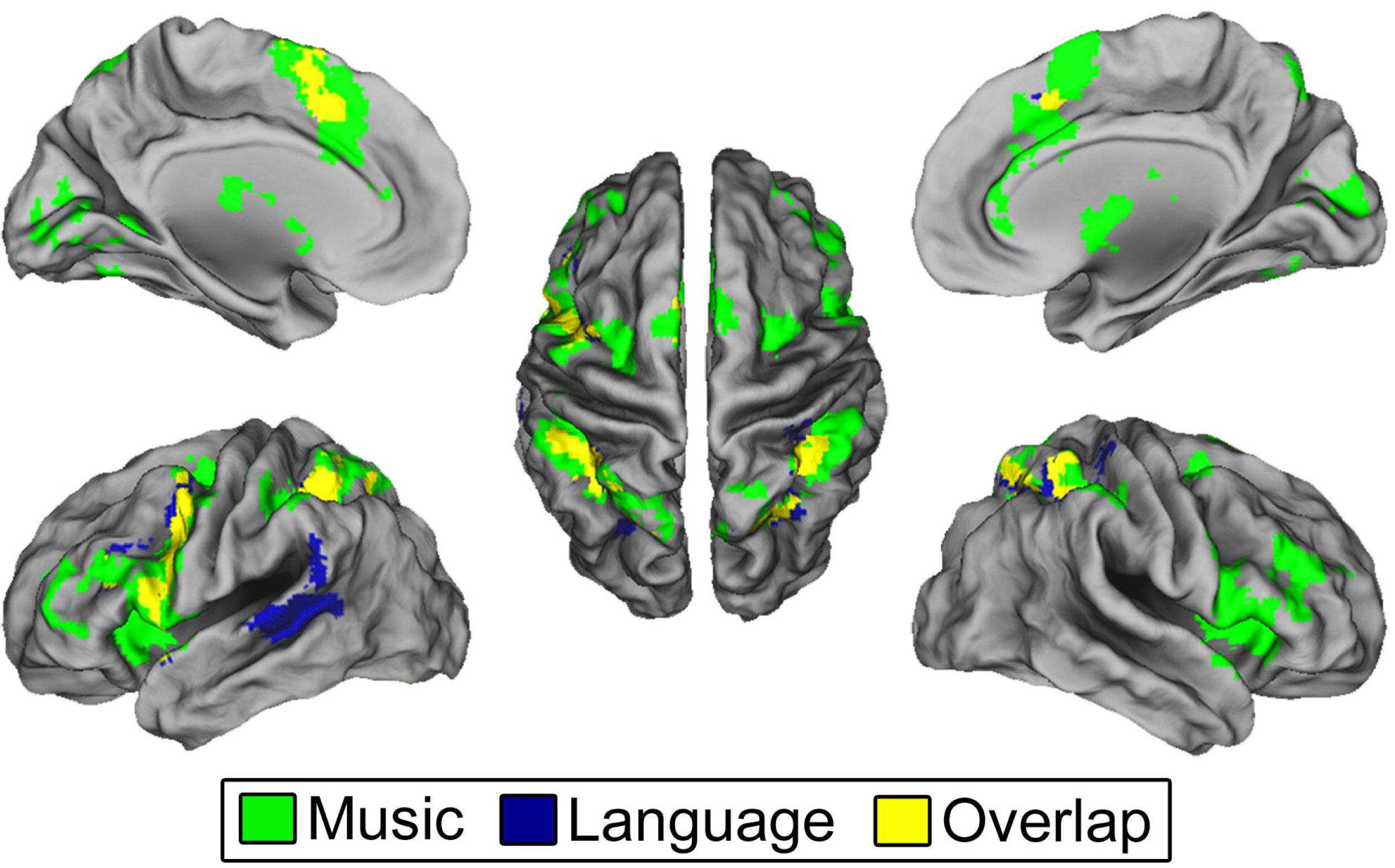
Group cross-classification (multivariate) result. Green regions represent areas in which the SVM algorithm could significantly classify, with above chance accuracy, structure vs. repeat trials in language materials after having been trained to recognize structure vs. repeat trials in music materials (i.e., M2L cross-classification). Blue regions represent areas in which the SVM algorithm could significantly classify, with above chance accuracy, structure vs. repeat trials in music materials after having been trained to recognize structure vs. repeat trials in language materials (i.e., L2M cross-classifications). Yellow areas show searchlight centers that can significantly perform both classifications.

As shown in Figure 3, significant cross-classifications were evident across a number of regions within medial prefrontal cortex, bilateral posterior parietal cortices, as well as left precentral, inferior (in the *pars opercularis*), and middle frontal gyri, matching areas of univariate overlap between the two domains (i.e., yellow regions in Figure 2). In addition, within each of these cross-classification clusters are areas (in yellow) capable of performing both L2M and M2L classifications, further demonstrating some extent of common underlying neural representation across the two domains. Crucially, this effect could be observed at the single-subject level, with median (single-subject) cross-classification accuracies at 61% for both L2M and M2L classifications, and ranges between 59% and 65%, and 58% and 64% for L2M and M2L classifications, respectively; with chance being 50%). Focusing on the left inferior frontal gyrus in particular, Figure 4 depicts the reliability of the result at the single-subject level. Figure 4 also demonstrates a significant across-subject variability in the exact location of voxels sensitive to linguistic structure within the inferior frontal gyrus (as previously shown; Fedorenko et al., 2010), something that we also observe in music-structure sensitive voxels, resulting in a systematic but variably located overlap in voxels capable of both L2M and M2L classifications within this region.

**Figure 4.**
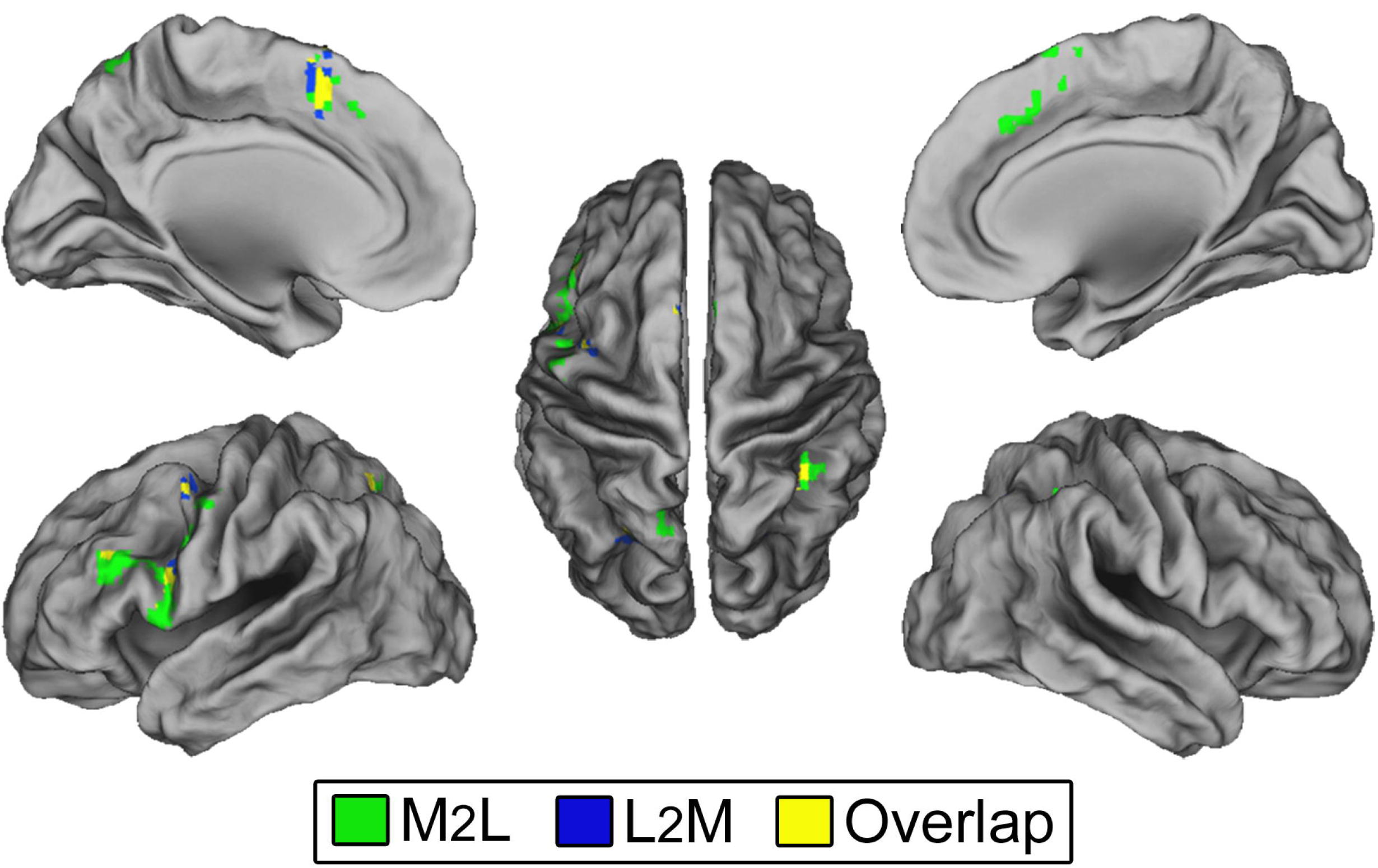
Single-subject cross-classification (multivariate) result. Cortical flat-maps depicting, for each participant separately, searchlight centers capable of significant cross-classifications (L2M in blue; M2L in green; overlap in yellow) within the inferior frontal gyrus as defined anatomically (highlighted in red, at the top). Each image (labeled as ‘S##’) represents the classification results for a single participant.

## Discussion

In this study we have addressed the question of the relationship between natural language and human cognition by contrasting the neural substrates accompanying the generation of structured sequences across language and music. Overall, our results provide the first direct evidence for the hypothesis that language and music have a shared neural code for producing structured relationships – a phenomenon that we observe both at the group as well as at the single subject level.

More specifically, we report three central findings. First, employing a magnitude-based univariate approach, we find the generation of structured sequences in language to recruit a well-known left lateralized network of frontal and temporal regions, along with posterior parietal foci, while generation of music sequences engaged a larger, and strongly bilateral, set of fronto-parietal regions. The neural substrate elicited by this performance paradigm (which has remained almost unexplored in the context of music, with the exception of Brown et al., 2006) matches very closely the neural substrate typically reported in tasks focusing on competence in both language (e.g., Ben-Shachar et al., 2003; Monti et al., 2009) and music (e.g., Maess et al., 2001; Koelsch et al., 2002, 2005).

Our second main finding, as evaluated with the same univariate activation-based approach, shows that the building of structured sequences in language and music relies on a number of common regions across left lateral and medial frontal cortices, as well as bilateral posterior parietal regions. In particular, the univariate analysis shows that the posterior-most aspect of Broca’s Area, in the *pars opercularis* of the left inferior frontal gyrus, is metabolically responsive to the presence of structure in the context of both language and music materials (Figure S2) – a finding that is consistent with results from previous studies (Maess et al., 2001; Koelsch et al., 2002, 2004, 2005; Brown et al., 2006). Beyond the left inferior frontal gyrus, our findings show that the interplay between language and music localizes to a set of regions in frontal and parietal cortices conventionally referred to as the multiple-demands network (Duncan, 2010), which have been shown to be recruited across a broad class of cognitive operations (Fedorenko et al., 2013). The absence of temporal regions identified by the domain-specific main effects (Figure S1), specifically the involvement of posterior STS (i.e. “Wernicke’s area”) in language tasks but not music, corroborates previous work reporting that posterior temporal regions might engage a semantic/syntactic interaction (See Friederici 2011 for a review).

Finally, our third, and crucial, finding addresses the significance of the frequently reported overlap between the neural substrate of language and that of music, thereby directly addressing the question of whether the mechanisms of natural language play a role in processing the structured sequences of music. Indeed, while regions of overlapping activation for these two domains have been widely interpreted as marking areas of shared neurocognitive processing (Kunert and Slevc, 2015). However, these hypotheses had not been directly tested (until now), prompting some to specifically advocate multivariate analyses such as the one adopted here (Peretz et al., 2015). As we reported above, we could find within each of the regions of univariate overlap (in Broca’s area), areas capable of recognizing music structure on the basis of language structure and vice versa. In fact, in each of these areas a subset of voxels could perform, at the same time, cross-classifications in both directions (i.e., L2M and M2L), demonstrating a degree of shared neuronal representation of structures across domains. These findings thus provide strong evidence in favor of the idea that language and music cognition share (neural) resources related to establishing structured relationships tying discrete elements into well-formed complex structures (Patel, 2003, 2012). Conversely, our results are at odds with the proposal that the overlap is either due to non-structural cognitive processes (e.g., salient events) or to the engagement of domain general regions proximal to Broca’s area (Fedorenko & Varley, 2016). Indeed, even at the single-subject level, regions sensitive to language and music structure appear to be variably positioned within the inferior frontal gyrus but systematically co-located (see Figure 4). Furthermore, the present findings also provide evidence in favor of the supramodal hierarchical parser hypothesis (Tettamanti & Weniger, 2006; Fadiga et al., 2009), according to which, encapsulated within Broca’s area, are mechanisms tied to processing hierarchical structure across multiple domains of human cognition. Nonetheless, whether the neural representation of specific operations (e.g., movement) can be directly mapped from one domain to another, as entailed by some views (Katz & Pesetsky, 2011), remains to be tested.

With respect to the idea that Broca’s area might serve as a supramodal parser of hierarchical structures, it is important to note, however, that to date it has only found support in a narrow sense (e.g., as conceived in Fadiga et al., 2009, and see Van de Cavey and Hartsuiker, 2015 for evidence of domain-general mechanisms), as it does not appear to extend to the hierarchical relationships of algebra (e.g., Varley et al., 2005; Monti et al., 2012), logic inference (e.g., Monti et al., 2007; Monti et al., 2009; Monti & Osherson, 2012), and spatial cognition (e.g., Bek et al., 2010). Thus far, it has only been found to be consistent with findings in the domain of language (cf., Bookheimer, 2002), music (here and in most previous neuroimaging reports; e.g., Maess et al., 2001; Koelsch et al., 2002; Koelsch et al, 2013), and motor action sequences (e.g., Fazio et al., 2009). In this sense, Broca’s area cannot be viewed as a central parser capable of operating in any domain of cognition as one would expect of a domain general processor or working memory component, though it may be a core component in a network of regions engaging in hierarchical processing (Fitch 2014). While it might be speculated that Broca’s area plays a role in cognitive domains where structured relationships trigger automatic and effortless intuitions (compare the ease of detecting a non-grammatical sentence or a sour note with the much more taxing task of detecting an incorrect algebraic expression or logic argument), the crucial factor(s) determining its involvement remains to be fully specified.

In conclusion, this report provides the first direct evidence that forging the structured sequences of natural language and music relies also on a common neural representational space which includes Broca’s area, a region traditionally associated with the syntactic operations of language. Whether the common mechanisms originally evolved in connection with one of the two domains, or whether they jointly inherited their properties from a common anteceding cognitive domain (e.g., a “prosodic protolanguage,” Fitch, 2005; or the capacity for hierarchical sequences of motor actions, Lashley, 1951), remains to be understood.

## Acknowledgments

This research was supported by the UCLA OVCR-COR Transdisciplinary Seed Grant “Language, Music, and the Brain” to A.L. and M.M.M., the National Defense Science and Engineering (NDSEG) fellowship to J.N.C., and by the Staglin IMHRO Center for Cognitive Neuroscience at UCLA.

## Author Contribution

MMM and AL developed the study concept and secured the funding. MMM, MHR, CAB, and DS devised the study design. MHR conducted behavioral testing and neuroimaging data collection. JNC and MHR performed the data analysis and, together with MMM, interpreted the results. MMM drafted the manuscript, MHR and JNC provided critical revisions. All authors contributed to subsequent editing of the manuscript.

